# Alien chromatin other than the GST-encoding *Fhb7* candidate confers Fusarium head blight resistance in wheat breeding

**DOI:** 10.1101/2021.02.03.429547

**Authors:** Xianrui Guo, Qinghua Shi, Jing Yuan, Jing Zhang, Mian Wang, Jing Wang, Chunhui Wang, Shulan Fu, Handong Su, Yang Liu, Yuhong Huang, Chang Liu, Qian Liu, Yishuang Sun, Long Wang, Ke Wang, Donglin Jing, Pingzhi Zhang, Jinbang Li, Houyang Kang, Yonghong Zhou, Xingguo Ye, Fangpu Han

## Abstract

The lack of resistance resources is a major bottleneck for wheat Fusarium head blight (FHB) resistance breeding. Three wheat-*Th. elongatum* FHB resistant translocation lines have been developed and used for wheat breeding without yield penalty. Transcriptomic analysis identified a derivative glutathione S-transferase transcript T26102, which was homologous to *Fhb7* and induced dramatically by *Fusarium graminearum*. Unlike other studies, *Fhb7* homologs were detected not only in *Thinopyrum* but also in *Elymus*, *Leymus*, *Pseudoroegeria* and *Roegeria*. We also found that several wheat-*Th. ponticum* derivatives carrying *Fhb7* and its homologs were highly susceptible to FHB. Moreover, the transgenic plants expressing *Fhb7* and its homolog on different backgrounds did not improve the FHB resistance.

**One Sentence Summary:** The GST-encoding *Fhb7* candidate cannot improve Fusarium head blight resistance in wheat breeding.

## Introduction

Fusarium head blight (FHB), colloquially known as scab, is caused by *Fusarium* species and remain one of the most devastating wheat diseases in many areas around the world (*1*). FHB causes significant yield losses and reduces grain quality because of mycotoxins generated during the course of infection, such as deoxynivalenol (DON) and nivalenol (NIV) (*2, 3*). Due to climate change, residue incorporation and maize/wheat rotations, FHB now occurs at greater and greater frequencies, making the task of developing FHB resistance wheat cultivars an urgent priority for breeders (*4–6*). While a few wheat accessions with high levels of FHB resistance have been exploited worldwide, most of them produce small heads, late maturity and other undesirable agronomic traits that hampered incorporating the resistance to elite cultivars (*3*). Cultivars Sumai 3 and its derivatives carrying *Fhb1* showed resistance to FHB and have been successfully applied to wheat breeding worldwide (*3, 7*). However, in spite of a successful clone of the major resistance gene *Fhb1*, it is difficult to combine the high FHB resistance with other necessary traits in wheat breeding practice, especially for winter and facultative wheat (*3, 7, 8*). It is of great significance to create novel FHB resistance resources that can combined with ecological adaptability and high yield.

The genus *Thinopyrum* contains numerous resistance genes for biotic and abiotic stress and is considered as an important genetic resource for wheat improvement (*9*). Until now, two genes conferring wheat FHB resistance were mapped to the homologous group seven in *Thinopyrum elongatum* and *Thinopyrum ponticum* (*10, 11*). Our previous research showed that the wheat-*Th. elongatum* 7E disomic addition line and substitution lines exhibit high resistance to FHB, and the further analysis confirmed the location of the resistant gene on the chromosome arm 7EL (*10*). Using a recombinant inbred line population from a cross between an FHB-susceptible substitution line (7E1/7D) and an FHB-resistant substitution line (7E2/7D) derived from *Th. ponticum*, the GST-encoding *Fhb7* candidate was reported to be located on the distal end of 7E2 and conferred broad resistance to *Fusarium* species by detoxifying trichothecenes via de-epoxidation (*11, 12*).

Many translocation lines have been developed from wild relatives of wheat and applied to wheat breeding. Some of the most successful example of transferring alien genes to common wheat include the wheat-rye 1BL/1RS translocation lines with fused centromere (*13*). These translocation lines were employed in wheat breeding because of their excellent stripe rust and powdery mildew resistance (*14*). Here, we produced a bountiful number of wheat-*Th. elongatum* translocation lines by radiating the pollens of the ditelosomic addition line 7EL and successfully applied the FHB resistant translocation lines to wheat breeding without yield penalty. In the process of screening the FHB resistance genes, we found the GST-encoding *Fhb7* candidate and its homologs are not responsible for FHB resistance.

## Results

### Development of wheat-*Th. elongatum* translocation lines

Previously, the long arm of chromosome 7E of *Th. elongatum* was found to harbor a new resistance gene capable of suppressing FHB spreading on wheat spikes (*10*). Irradiation was performed on the pollens of wheat-*Th. elongatum* addition line 7EL at early flowering stage. Following that, fresh pollens were pollinated to the emasculated spikes of the recurrent parent. In total, 8400 M1 seeds were obtained and cytological analyses were performed on all germinated seeds. Consequently, 671 wheat-*Th. elongatum* translocation lines were identified, accounting for 7.99% of all developed lines (Table 1). The translocation lines were classified into terminal and intercalary types by the position of alien chromosome fragments (fig. S1A-D). By the size of alien chromosome fragments, the terminal translocation lines were furtherly classified into short, medium and long alien segmental translocation lines (fig. S1A-C). In total, we obtained 184 short alien segmental translocation lines, 141 medium alien segmental translocation lines, 247 long alien segmental translocation lines and 99 intercalary translocation lines (Table 1).

**Table 1.** Four types of translocation lines identified from 8400 seeds.

| Translocation type | Number | Percentage (%) |
| --- | --- | --- |
| Short alien segmental translocation line | 184 | 27.42 |
| Medium alien segmental translocation line | 141 | 21.01 |
| Long alien segmental translocation line | 247 | 36.81 |
| Intercalary translocation line | 99 | 14.76 |
| In total | 671 | 100 |

### Application of wheat-*Th. elongatum* translocation lines in FHB resistance breeding

Wheat cultivar Jimai 22 is widely grown in northern China because of its broad adaptation and high yield potential. In order to improve the integrated agronomic traits of our developed translocation lines, we back crossed them with Jimai 22 and evaluated FHB resistance using the single floret inoculation method. After one to three generations of backcross 81 translocation lines were identified with good resistance to FHB (Fig.1A-C and fig. S2). Karyotype analysis showed that translocations occurred among all seven homologous groups in wheat (fig. S2). After backcrossing with Jimai 22 for at least three generations, homozygous translocation lines were selected from the self-crossed progenies. Out of these, the short alien segmental translocation lines Zhongke 1878 and Zhongke 166 as well as the long alien segmental translocation line Zhongke 545 were developed with good integrated agronomic traits and no significant grain yield penalty (Fig. 1D and fig. S3). All the three lines showed high resistance to FHB, with less diseased spikes and less kernels than the control cultivar Jimai 22 after infection in nature (Fig. 1E and F). Field assessment across China in a national contest showed that the grain yield of Zhongke 166 exceeded that of the national control Zhoumai 18 by an average of 6.26% in 23 locations (Fig. 1G). Cytological analysis revealed that the translocation occurred on the long arm of chromosome 6D in line Zhongke1878, and on the long arm of chromosome 7D in line Zhongke 166 and Zhongke 545 (fig. S3).

**Fig. 1.**
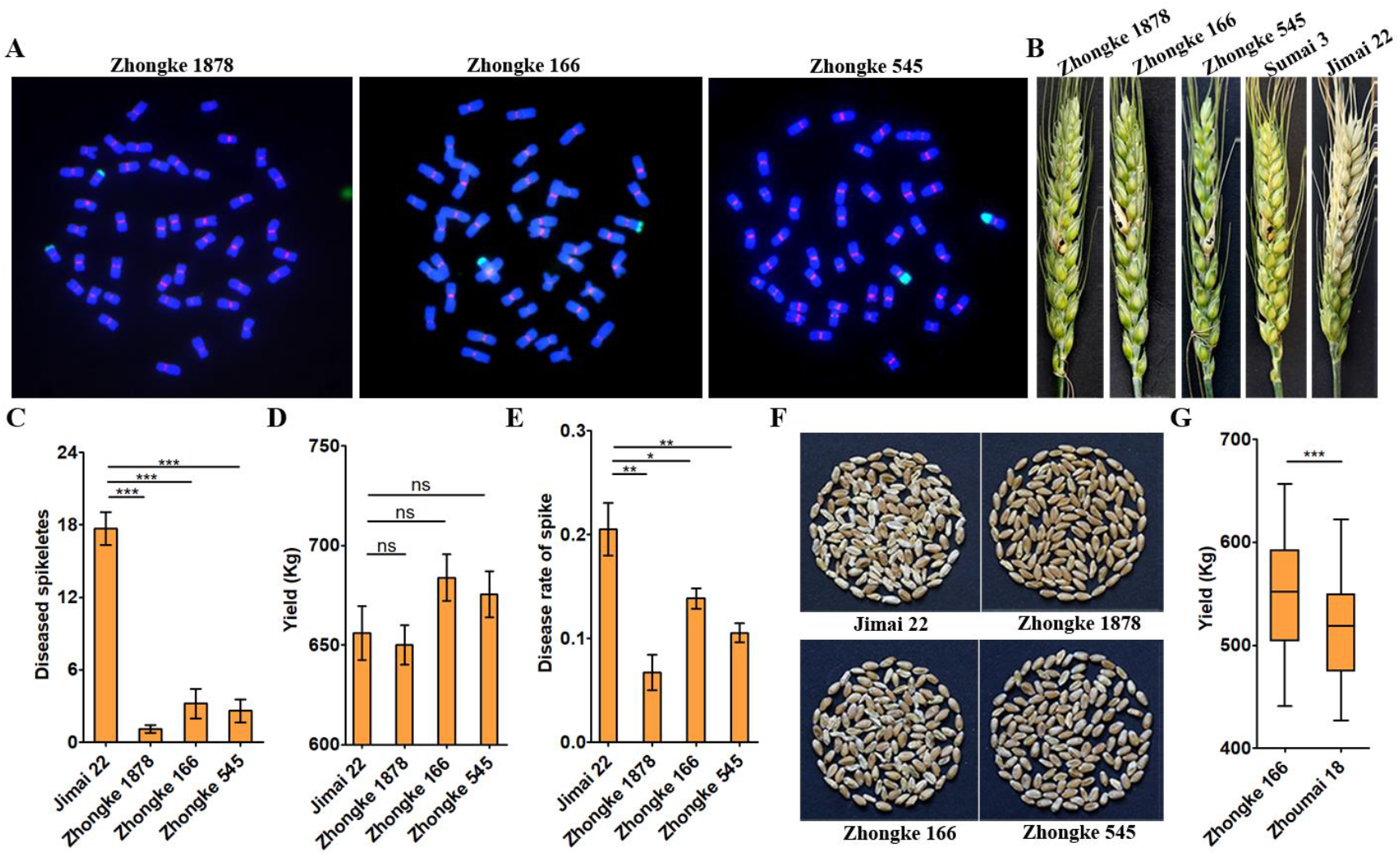
Three wheat-*Th. elongatum* translocation lines with FHB resistance and yield advantage. (A) FISH analysis of the translocation lines Zhongke 1878, Zhongke 166 and Zhongke 545. The green signal identifies *Th. elongatum* chromatin in wheat background. The red signal represents wheat centromere. DAPI stains chromosome blue. (B-C) FHB resistance evaluation on three wheat-*Th. elongatum* translocation lines. The diseased spikes were photographed (B) and the number of diseased spikeletes were calculated (C) at 21 d after inoculation with *Fusarium graminearum*. (D) Grain yield of three translocation lines in plot scale in Beijing. All translocation lines and the control Jimai 22 were grown in nature field and the area of the trial plot is 13.3 m^2^. (E-F) The diseased rate of spike (E) and diseased kernel (F) under nature field condition in Anqing. (G) Yield comparison between the translocation line Zhongke 166 and the national control Zhoumai 18. The yield data were collected from the national wheat yield contest across 23 locations. The statistics analysis were conducted by t test (*P < 0.05, **P < 0.01, ***P < 0.001).

### Screening specific transcripts for disease-resistant interval on chromosome 7EL

To explore the nature of the FHB resistance gene in 7EL, we inoculated the spikes of translocation line Zhongke 1878 with *Fusarium graminearum* and performed full-length transcriptome sequencing after ninety-six hours. Among 34996 transcripts, 520 transcripts were identified as candidates expressing only from 7EL by blasting against the reference genome of Chinese Spring (CS) and the nucleotide database of *Fusarium graminearum* on the National Center for Biotechnology Information (NCBI). According to the identified sequences, one to three pairs of primers for each transcript were designed. In order to identify genes specifically located on the alien chromatin from 7EL, polymerase chains reaction (PCR) was performed using genomic DNA of wheat-*Th. elongatum* addition line 7EL, CS, Zhongke 1878 and Jimai 22. Finally, 25 genes specific to the disease-resistant interval in line Zhongke 1878 were obtained (fig. S4A and data S1). Among them, 7 transcripts were annotated as resistant proteins containing the NB-ARC domain and 10 as unknown proteins (Table 2). Additionally, other proteins found included proteins such as a receptor kinase, ATPase subunit, dirigent-jacalin protein, GST family, nucleosome assembly protein, cold induced protein and retrotransposon protein (Table 2). Annotated as a GST protein, T26102 was chosen for further study because it was induced drastically 48h after inoculation with *Fusarium graminearum*, which was confirmed by qRT-PCR (Fig. 2A and fig. S4B). We also collected the RNA-seq data of CS-7EL line 4 d after water and *Fusarium graminearum* inoculation previously reported from NCBI (*15*), which further confirmed the induction of T26102 by *Fusarium graminearum* (fig. S4C).

**Table 2.** The transcripts specific for disease-resistant interval in line Zhongke 1878.

| Transcripts | Length (bp) | Function annotation |
| --- | --- | --- |
| T255 | 5439 | CC-NBS-LRR protein |
| T1375 | 4353 | CC-NBS-LRR protein |
| T2475 | 4015 | unknown |
| T2921 | 3963 | NB-ARC domain |
| T4619 | 3607 | unknown |
| T6683 | 3480 | unknown |
| T6981 | 3503 | CC-NBS-LRR protein |
| T7667 | 3443 | CC-NBS-LRR protein |
| T8538 | 3325 | unknown |
| T9007 | 3358 | CC-NBS-LRR protein |
| T10410 | 3204 | unknown |
| T11006 | 3228 | CC-NB-ARC domain |
| T14242 | 2493 | Retrotransposon protein |
| T15795 | 2233 | Lectin receptor kinase |
| T19432 | 1873 | DNA repair exonuclease SbcCD |
| T19609 | 1861 | unknown |
| T20934 | 1769 | Retrotransposon protein |
| T26102 | 1394 | GST protein |
| T26788 | 1389 | Dirigent-Jacalin protein |
| T28749 | 1249 | Nucleosome assembly protein |
| T30581 | 1098 | unknown |
| T32029 | 996 | unknown |
| T32111 | 993 | unknown |
| T32749 | 934 | Freezing-induced protein |
| T33754 | 844 | unknown |

**Fig. 2.**
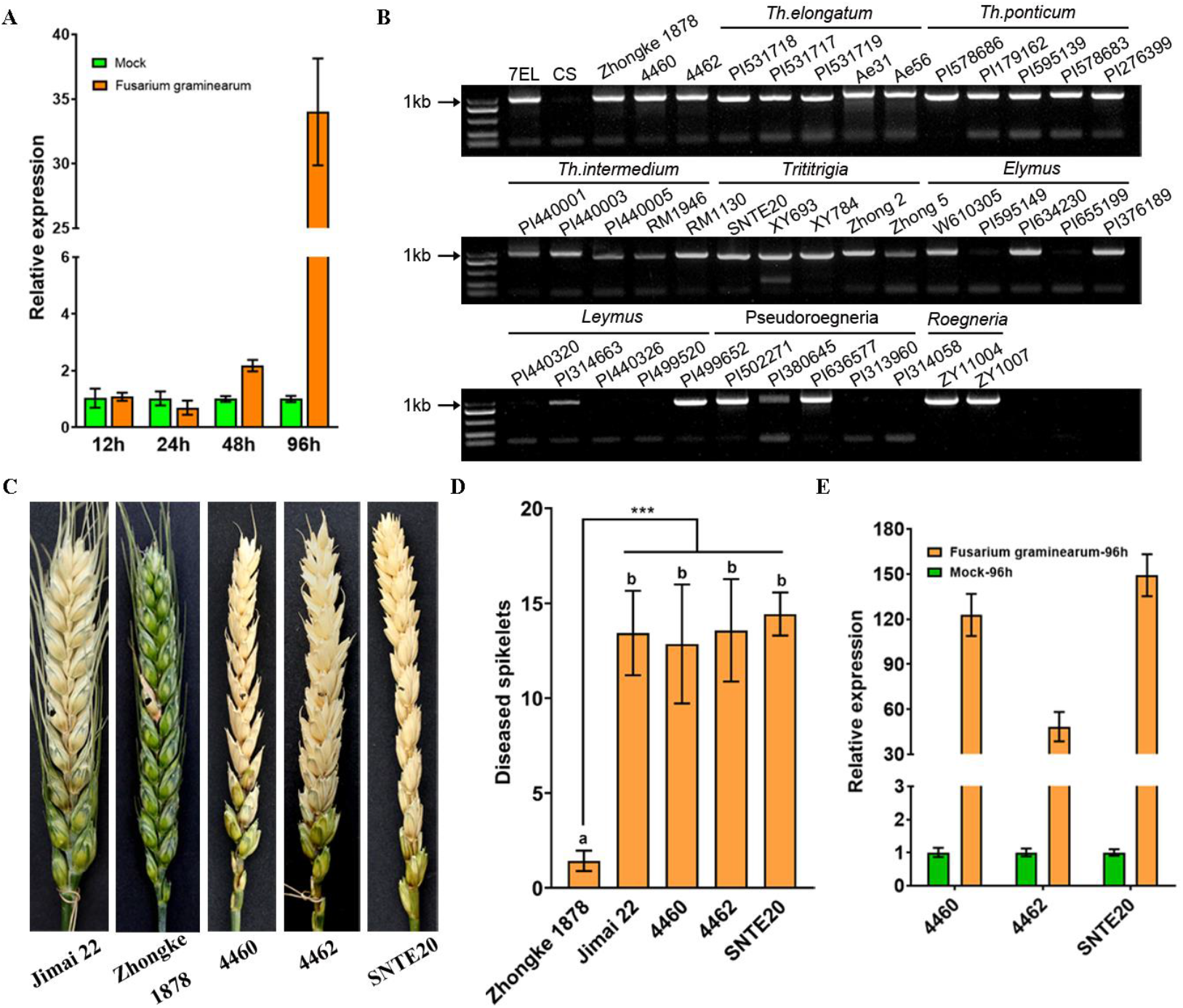
The expression pattern and FHB resistance evaluation on some amphiploids and translocation lines carrying the *Fhb7* homolog. (A) Expression pattern of T26102 in Zhongke 1878 after inoculating with *Fusarium graminearum*. The Mung bean soup without *Fusarium graminearum* was used as mock treatment. (B) Detection of the T26102 homolog among different species in Triticeae. (C-D) FHB resistance evaluation on wheat-*Th. ponticum* translocation lines and amphiploid. The diseased spikes were photographed (C) and the number of diseased spikeletes were calculated (D) at 14 d after inoculation with *Fusarium graminearum*. Zhongke 1878 was used as resistant control and Jimai 22 as susceptible control. 4460 and 4462, wheat-*Th. ponticum* translocation lines; SNTE20, wheat-*Th. ponticum* partial amphiploid. (E) Expression comparison of the *Fhb7* homolog among 4460, 4462 and SNTE20. The statistics analysis were conducted by t test (***P < 0.001).

### Distribution of T26102 in Triticeae

To explore the evolution of T26102, its homologs were checked in different wheat-*Thinopyrum* derivates. The T26102 homologs were detected in the addition line 7EL, our developed wheat-*Th. elongatum* translocation lines, such as Zhongke 1878, and the wheat-*Th.ponticum* translocation lines 4460 and 4462 (Fig. 2B). The homologs of T26102 were also detected in wheat-*Thinopyrum* partial amphiploids, such as octoploid SNTE20 and XY693 derived from the hybridization between hexaploid wheat and *Th. ponticum*. Totally, 122 accessions belonging to Triticeae were collected and used for detecting the T26102 homologs (data S2). Except for *Thinopyrum*, the homologs of T26102 were detected in four other genera, i.e., *Elymus*, *Leymus*, *Pseudoroegneria* and *Roegneria* (Fig. 2B).

By comparing sequences, we found that T26102 was homologous to the reported *Fhb7* with only two amino acids difference between them. Furthermore, the protein sequences were at least 95% identical across all the *Fhb7* homologs in Triticeae plants (fig. S5). In some species, more than one homolog was discovered. We detected two homologs of *Fhb7* in our developed wheat-*Th. elongatum* translocation lines, such as Zhongke 1878 and Zhongke 166 (fig. S5). Three homologs of T26102 were detected in the *Th. intermedium* accession PI 440001 (fig. S5). Despite indel variation and amino acid substitution across all the homologs of *Fhb7*, no premature termination and code-shifting mutations occurred in the protein sequences. The main variation was the number of Thr-Ser at the amino terminus of the protein sequence (fig. S5).

### Functional identification of*Fhb7* homologs

To identify the function of the GST-encoding *Fhb7*, we evaluated the FHB resistance on wheat-*Thinopyrum* derivatives carrying the homologs of *Fhb7*. Surprisingly, obvious differences in FHB resistance were detected among different lines carrying the homolog of *Fhb7*. The wheat-*Th. ponticum* translocation lines 4460, 4462, and wheat-*Th. elongatum* translocation line Zhongke 1878 all carried the *Fhb7* homolog (Fig. 2B and fig. S6). However, the translocation lines 4460 and 4462 were susceptible to FHB, whereas the Zhongke 1878 was resistant to FHB (Fig. 2C and D). Sequence alignment analysis revealed that the protein sequences in 4460 and 4462 lines were identical to the reported *Fhb7* from the 7E2/7D substitution line (fig. S5). Furthermore, one Wheat-*Thinopyrum* partial amphiploids carrying the *Fhb7* homolog were also identified as susceptible to FHB, such as octaploid SNTE20 (Fig. 2B-D and fig. S6). Expression analysis revealed that the expression of *Fhb7* homologs was induced in 4460, 4462 and SNTE20 after inoculating with *Fusarium graminearum* (Fig. 2E). We also discovered that lines 4460 and 4462 shared an identical promoter with *Fhb7* from the 7E2/7D substitution (fig. S7). These results thus raised serious questions about the ability of the GST-encoding *Fhb7* to confer resistance to FHB.

To verify the FHB resistance function of T26102, we transformed the overexpression vector pUbi:T26102 into three wheat accessions 19AS161, Jimai 22 and Zhongmai 175, all of which are highly susceptible to FHB. The transgenic positive wheat plants overexpressing T26102 were used for FHB resistance evaluation (Fig. 3A and fig. S8A). Compared to the wild types, no T0 transgenic lines showed an improved FHB resistance regardless of any genetic background (Fig. 3B and C). This result further suggests that T26102 is not the pivotal gene that confers wheat with resistance to FHB in our translocation lines. To rule out the effect of amino acid variation on the function of T26102, we also expressed the GST-encoding *Fhb7* under the *ubiquitin* promoter and the native promoter on common wheat varieties Zhengmai 7698 and Kenong 199. The transgenic positive wheat plants expressing the *Fhb7* were used for FHB resistance evaluation (fig. S8B-D). As with the control, the T0 generation transgenic plants on Zhengmai 7698 background and the T1 generation transgenic plants on Kenong 199 background were identified as highly susceptible to FHB (Fig. 3D and E). Regardless of the *ubiquitin* or the native promoter, there is no difference between the T1 transgenic plants and the control Kenong 199 (Fig. 3F).

**Fig. 3.**
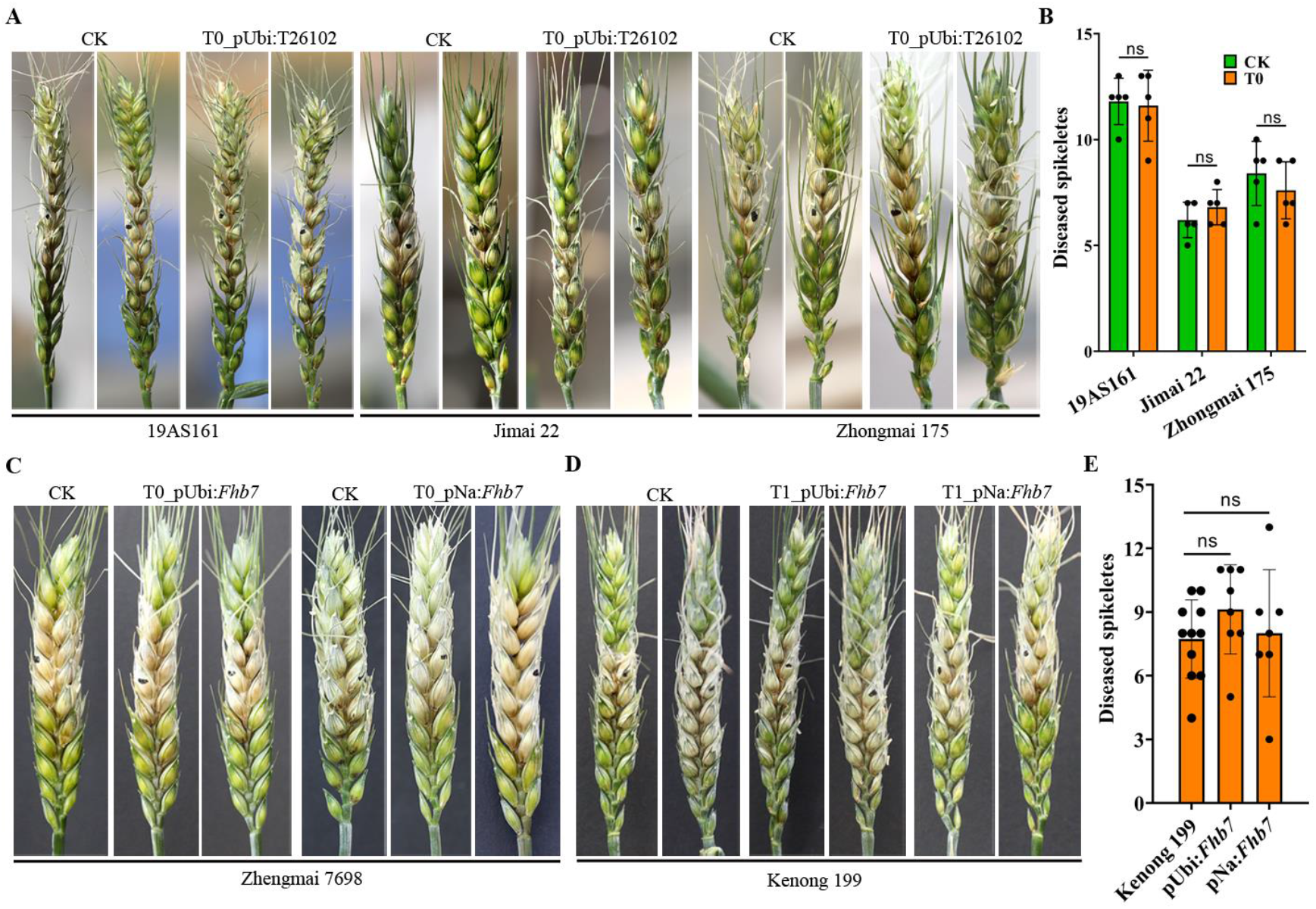
FHB resistance evaluation on the transgenic lines with *Fhb7* homolog. (A-B) FHB resistance evaluation on the control (CK) and T0 transgenic lines overexpressing T26102. The diseased spikes were photographed (A) and the number of diseased spikeletes were calculated (B) at 7 d after inoculation with *Fusarium graminearum*. (C) FHB resistance evaluation on the T0 transgenic lines expressing *Fhb7* on the background Zhengmai 7698. pUbi:*Fhb7* indicated that *Fhb7* was driven by the *ubiquitin* promoter. pNa:*Fhb7* indicated that *Fhb7* was driven by the native promoter. The diseased spikes were photographed at 7 d after inoculation with *Fusarium graminearum*. (D-E) FHB resistance evaluation on the T1 transgenic lines expressing *Fhb7* on the background Kenong 199. The diseased spikes were photographed (D) and the number of diseased spikeletes were calculated (E) at 7 d after inoculation with *Fusarium graminearum*.

## Discussion

Crop wild relatives are undeniably beneficial to modern agriculture because they provide breeders with a broad pool of potentially useful genetic resources, especially with regard to the resistance to disease and pest (*16*). Translocation lines between wheat and its wild relatives have been successfully applied to wheat breeding, such as the wheat-rye 1BL/1RS translocation line and the wheat-*Haynaldia villosa* 6AL/6VS translocation line (*13, 17*). The usefulness of the translocation lines is dependent on whether the alien fragment could compensate for the replaced wheat segments (*18*). In our study, translocation lines between wheat and *Th. elongatum* were successfully applied to wheat FHB resistance breeding without yield penalty, and the translocation for Zhongke 166 and Zhongke 545 occurred on chromosome 7DL. Their high yield potential in the pilot experiment might be attributed to the compensation from the translocated 7EL segment with the lost 7DL segment. Our practice revealed that Zhongke 1878, whose translocation occurred on 6DL, also exhibited good yield potential. This result suggests that the application of translocation lines should not be limited to the translocations between homeologous chromosomes.

It is well known that germplasm from East Asia harbors highly resistant genotypes. Chinese cultivar Sumai 3 and its related lines have been used in breeding project worldwide because of their high resistance to FHB (*19*). It is evident that most of the Sumai 3 derived resistant cultivars released in the United States, Canada and China belong to the spring type (*20, 21*). Apart from the French cultivar Jaceo (carrier of *Fhb1*) which was on the market only for a short period, no further winter wheat cultivars harboring *Fhb1* have been released so far because of poor adaptation traits (*20, 21*). Therefore, there is significant urgency in breeding practice to incorporate FHB resistance into winter varieties. In our breeding practice, we successfully incorporated the alien chromatin with FHB resistance into the major sowing varieties Jimai 22 with no sacrifice of yield or adaptability. Therefore, the translocation lines Zhongke 1878, Zhongke 545 and Zhongke 166 could be used as novel FHB resistant resources for wheat improvement in China, especially for the Yellow and Huai River Valleys that relies on winter and facultative wheat types. In order to further improve FHB resistance, we plan to incorporate the alien resistant chromatin into the “Ningmai” and “Yangmai” lines, which play an important role in the Middle and Lower Yangtze Valleys.

Small segmental translocation lines with Fusarium head blight resistance can help narrow down the region carrying the resistant gene. We applied this strategy to our work, and selected twenty-five transcripts specific for disease-resistant regions on 7EL. We focused on T26102 because of its dramatical induction after inoculation with *Fusarium graminearum* and its similarity to *Fhb7*. Recently, *Fhb7*, also encoding the GST protein, was reported to confer wheat with a broad resistance to *Fusarium* species and it was acquired by horizontal gene transfer (HGT) from *Epichloë* to *Thinopyrum* (*11*). Interestingly, except for *Thinopyrum*, we were able to find homologs of *Fhb7* in several species within the genera of *Elymus*, *Leymus*, *Pseudoroegeria* and *Roegeria*. This was unsurprising, given that *Epichloë* often formed symbiotic associations with temperate grasses of the subfamily *Pooideae* (*22*). Thus to us HGT did not appear to be an accidental happening by chance only in *Thinopyrum*. It is possible that HGT happened before Triticeae differentiation, or else *Fhb7* ought to be detected in the genus *Triticum* at large, which currently does not appear to be the case.

Perhaps more interesting is the fact that the GST-encoding *Fhb7* and its homologs are not causal in conferring FHB resistance. Regardless of *Fhb7* or its homolog T26102, none of the positive transgenic lines was able to improve FHB resistance. Furthermore, some wheat-*Thinopyrum* translocation lines and partial amphiploids carrying the homologs of T26102 were also identified as susceptible to FHB. Two *Fhb7* homologs were detected in the wheat-*Th. ponticum* amphiploid SNTE20 and their protein sequence was 98% homologous to the *Fhb7* cloned from *Th. elongatum*. However, SNTE20 was identified as highly susceptible to FHB. Above all, wheat-*Th. ponticum* translocation lines 4460 and 4462, that carried an identical protein sequence to Fhb7 in the 7E2/7D substitution line also failed to confer FHB resistance in our experiment. These results indicate to us that T26102, including its homolog *Fhb7*, are not responsible for FHB resistance.

Studies based on meiotic chromosome pairing revealed that *Th. elongatum* chromosome 7E paired occasionally with *Th. ponticum* chromosomes 7E1 and 7E2 in hybrids, with frequencies of meiotic pairing rates of 19.85 and 2.52, respectively (*23*). The genetic relationships based on molecular markers also revealed that 7E was distant from 7E1 and 7E2 (*23*). It was reported that resynthesized polyploids and natural polyploids have undergone many genetic changes, including sequence deletion, rDNA loci changes, transposon activation and chromosomal rearrangement (*24–30*). All these findings suggest that 7E and 7E2 have difference on the DNA level. *Fhb7* was mapped to the distal end of 7E2 between the XSdauK79 and XSdauK80 markers based on recombinations between 7E1 and 7E2 (*11, 23*). Although ~1.2 Mb region between the two markers was identified on the 7E chromosome, the DNA components between the two mapping intervals of 7E and 7E2 might be quite different. The fortuitous findings of FHB resistant candidate genes from *Th. ponticum* are unlikely to be replicated using *Th. elongatum* as a reference genome.

The lack of resistance conferred by the *Fhb7* and its homolog T26102 in our work indicates that the bona fide resistance gene still lies undiscovered. So long as FHB remain as a major threat to worldwide agriculture, identifying and exploiting its resistance genes will always prove to be a useful endeavor

## Supporting information

The sequences of twenty-five specific transcripts in line Zhongke 1878

Distribution of the Fhb7 homologs in Triticeae

## Acknowledgments

We thank X.E. Wang (Nanjing Agricultural University) and J.B. Li (Nanyang Academy of Agricultural Sciences) for providing the *Fusarium* strains, Y.G Bao and H.G. Wang (Shandong Agricultural University) for providing some partial amphiploid accessions, Y.W. Li (Institute of Genetics and Developmental Biology, Chinese Academy of Sciences) and Moshe Feldman (Weizmann Institute of Sciences) for providing *Thinopyrum* species, and M.C. Luo (UC Davis Plant Sciences) for providing the wheat-*Th. elonatum* addition line 7EL.

## Funding

This work was supported by the National Key Research and Development Program of China (2016YFD0102001).

## Author contributions

F.H. designed the project. X.G., M.W. and J.Y. conducted the experiment and performed data analysis. F.H., Q.S., J.Z., J.W., C.W., L.W., H.S., Y.L., Y.H., C.L., Q.L., Y.S., S.F., D.J., P.Z., J.L and Y.Z conducted the field work. K.W. performed the wheat genetic transformation. X.G., J.Y., X.Y and F.H. wrote the paper.

## Competing interests

The authors declare no competing interests.

## Data and materials availability

All data are available in the manuscript, the supplementary materials or at the publicly accessible repositories. These data in the public repositories include all transcriptomic raw reads for the translocation line Zhongke 1878 in NCBI under BioProjectID PRJNA720120. All materials were available from Fangpu Han.

## Materials and Methods

### Plant materials

All 122 species belonging to Triticeae were listed in Supplementary Data 2. The wheat-*Thinopyrum ponticum* translocation lines 4460 and 4462 were obtained from Agriculture and Agri-Food Canada (provided by George Fedak). Octoploid trititrigia SNTE20, SNTE122, XY693, XY784 and XY7631 were produced by hybridization between *Thinopyrun ponticum* and common wheat (provided by Honggang Wang and Zhensheng Li). The accession 19AS161 with high transformability were produced from CB037/Kenong 199//Kenong 199///Kenong 199. CB037 was a tissue-culture-favorable common wheat line provided by Xingguo Ye (*31*). The released common wheat varieties Jimai 22, Zhongmai 175 and Zhengmai 7698 were highly susceptible to FHB and used as the transgenic receptors.

### Induction and improvement of wheat-*Th. elongatum* translocation lines

At flowering stage, the spikes of wheat-*Th. elongatum* addition line 7EL were cut from the plant in the morning, and immediately radiated by γ rays derived from ^60^Co with a dose of 18 Gy. Then the fresh pollens were pollinated to the cultivated variety Jimai 22 with the stamens removed in advance. The hybrid seeds were harvested at maturity. The translocation lines were identified by utilizing fluorescence *in situ* hybridization (FISH). The translocation lines in which the ratio of the length of the alien fragment to the full length of 7EL ranged from 0 to 1/4 were considered to be short alien segmental types. The translocation lines in which the ratio ranged from 1/4 to 1/2 or from 1/2 to 1 were classified as medium or long alien segmental types. Using Jimai 22 as the recurrent parent, the integrated agronomic traits of the translocation lines were improved by continuous backcrossing.

### Fluorescence*in situ* hybridization (FISH)

The translocation lines were screened by FISH according to previously reported methods (*25*). The seeds harvested were germinated on moist filter paper in a petri dish at room temperature for 2-3 d. The roots were cut from the seedlings and then placed in nitrous oxide for 2h. Subsequently the roots were fixed in 90% acetic acid for 5 minutes and then washed three times by sterile water. Chromosome spreads preparation was performed as previously described (*32*). 7EL-1 was obtained by Dop-PCR from the 7EL library constructed by chromosome microdissection. It was specific for the genome of *Th. elongatum* and *Th. ponticum*. The probes were labeled using the nick translation method (*33*). Two repetitive sequences pAsⅠ and pSc119.2 were used to identify the whole set of wheat chromosomes. 7EL-1 and pSc119.2 were labelled with Alexa Fluor-488-5-dUTP. The centromeric retrotransposon of wheat clone 6C6 and pAsⅠ were labeled with Texas-red-5-dCTP.

### Fusarium head blight resistance evaluation

During the year 2015-2017 in field condition in Beijing (E116°42′, N40°10′), FHB evaluations were performed by using the single spikelet inoculation method (*34*). Equally mixing three pathogenic *F. graminearum* strains (Fg16-2, Fg16-5 and Fg16-11) and one *Fusarium asiaticum* strain (Fa301) in Mung bean broth produced fungal spores. For convenience, we referred to the four mixed species as *Fusarium graminearum*. Approximately 20 μL of *F. graminearum* fungal suspension (1× 10^6^ conidia/ml) was injected into the central spikelet at early flowering stage. For each wheat-*Th. elongatum* translocation lines, at least 10 spikes were inoculated with *Fusarium graminearum* each year. The recurrent parent Jimai 22 was used as susceptible control and was planted by stages. For the transgenic plants, at least three positive individual plants were used for FHB resistance evaluation. The inoculated spikes were covered with a plastic bag for 2 d to keep moist for fungal infection. The percentage of diseased spikelets for each spike was recorded at 7 d, 10 d and 21 d after inoculation. For the translocation lines Zhongke 1878, Zhongke 166 and Zhongke 545, the diseased rate of spikes were also calculated and repeated three times in nature condition in Anqing (E117°14′, N30°64′) in the year 2019. The diseased rate of spikes were statisticted at 21d after flowering. The picture of diseased kernels were photographed after the translocation lines were harvested. The statistical analysis was performed by the unpaired t test using the software GraphPad Prism 8.

### Yield comparison experiment

For the translocation lines Zhongke 1878, Zhongke 545 and Zhongke 166, the yield comparison experiments were performed on a 13.3 m^2^ plot and were repeated three times in Beijing (E116°42′, N40°10 ′) during the year 2018-2019. The recurrent parent Jimai 22 was used as yield control. In the year 2020, the translocation line Zhongke 166 participate in the national regional trials named South of Huanghuai Area across 23 locations. The yield data was collected and compared with the national regional trial control wheat variety Zhoumai 18 released in the year 2005. The statistical analysis was performed by the paired or unpaired t test using the software GraphPad Prism 8.

### RNA sequencing and screening transcripts induced by *Fusarium graminearum*

To explore the resistance gene for Fusarium head blight, the spikes of the translocation line Zhongke 1878 were sampled for RNA sequencing after inoculating with *Fusarium graminearum*. Three spikelets around the inoculated one from at least three spikes of different plants were collected at 12, 24, 48, and 96h post inoculation and grounded in liquid nitrogen for total RNA extraction using TRIzol^®^ Reagent (Invitrogen). As lesions were observed on the glumes at 96h post inoculation, the sample at 96h was selected for full-length transcriptome sequencing. Firstly, we aligned the sequenced transcripts on the Chinese Spring reference genome by using the software GMAP (with parameters: -min-trimmed-coverage 0.9 -min-identity 0.85). To remove the transcripts derived from the inoculated *Fusarium graminearum* isolates, the unmapped transcripts were blasted against the nucleotide database on the National Center for Biotechnology Information (NCBI). In order to confirm their origin, one to three pairs of primers were designed for the transcripts left. Polymerase chain reactions were carried out using the genome DNA of wheat-*Th. elongatum* 7EL, Chinese Spring, Zhongke 1878 and Jimai 22. The functions of the transcripts specific for Zhongke 1878 were annotated by using Blastx on the NCBI. The data analysis was performed by employing HISAT2 and StringTie according to the previously report (*35*). The FPKM value from StringTie was used to measure the expression level.

### Distribution, expression analysis and genetic transformation of T26102

In order to detect the distribution of T26102 in different species in Triticeae, the fragment of T26102 was amplified by using the primer set of F-CGATAGAAGATAGCTTCAATCAACCCTTT and R-CTACTTCACCTCGGCATACTTGTC. The fragments amplified from different species were cloned onto the *pEASY*^○^R -T1 simple cloning vector (TransGen Biotech Co, Beijing) for sequencing. The sequence comparison analysis was carried out using the software DNAMAN. First-strand cDNA synthesis from the total RNA was performed by using the FastKing RT kit (with gDNase) (TianGen Biotech Co, Beijing). The expression analyses were performed using the primer set (F-GGACTTCCCTTGGATCCTGC and R-ACCGACAATCATGTCCGCAT). The gene *actin* was used as an internal standard by the primer set of F-CAACGAGCTCCGTGTCGCA and R-GAGGAAGCGTGTATCCCTCATAG. The relative expression of T26102 was calculated by the 2 ^−ΔΔCT^ method.

The 846 bp CDS of T26102 was amplified from the genomic DNA of the translocation line Zhongke1878. The CDS was cloned into the MCS of the modified pWMB110 vector under the *ubiquitin* promoter by using the EasyGeno Assembly Cloning kit (TianGen Biotech Co, Beijing). The recombinant plasmid was transformed into *Agrobacterium* strain C58C1 (Zoman Biotech Co, Beijing). The *Agrobacterium*-mediated wheat transformation using the immature embryos of 19AS161, Zhongmai 175, Jimai 22, Zhengmai 7698 and Kenong 199 was carried out as previously described (*36*). The positive T0 and T1 plants expressing T26102 confirmed by RT-PCR were used for FHB resistance evaluation.

## Supplementary Materials

**Fig. S1.**
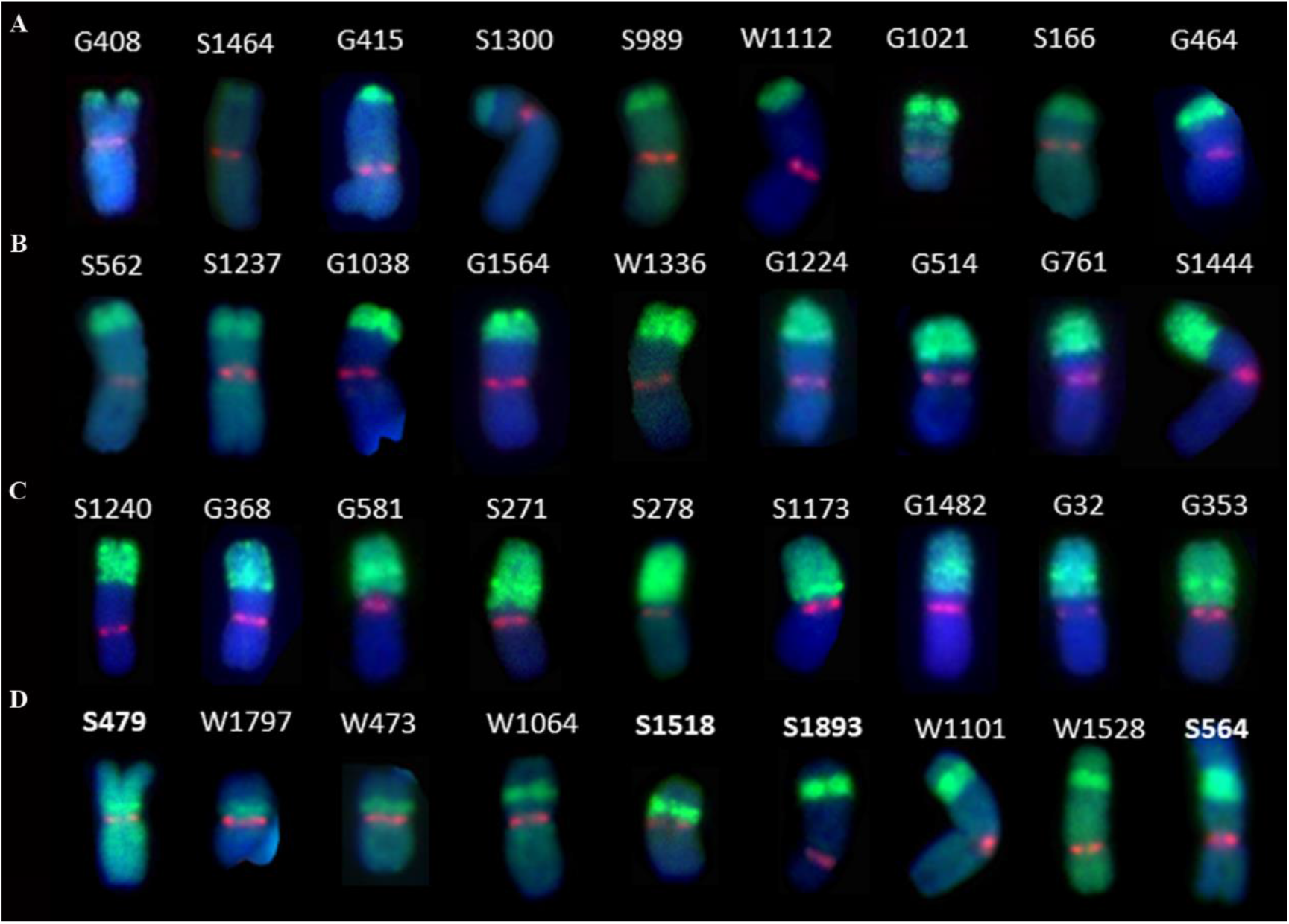
Four types translocation lines identified by FISH. (A) Short alien segmental translocation line examples. (B) Medium alien segmental translocation line examples. (C) Long alien segmental translocation line examples. (D) Intercalary translocation line examples. The green signal identifies *Th. elongatum* chromatin in wheat background. The red signal represents chromosome centromere. DAPI stains chromosome blue.

**Fig. S2.**
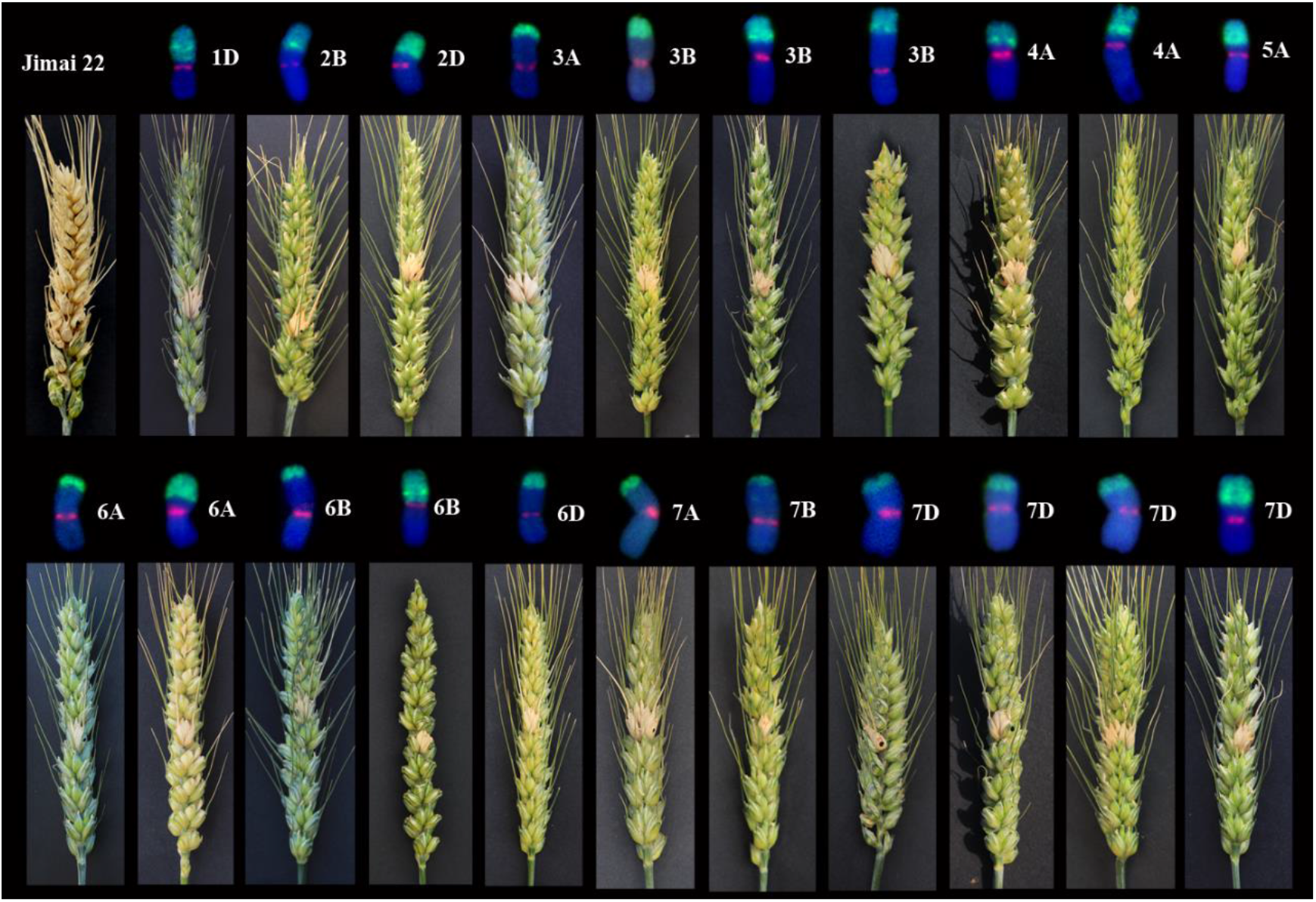
FHB resistant translocation lines identified by single spikelet inoculation method. Except for the susceptible control Jimai 22, each spike represent one wheat-*Th. elongatum* translocation line. The translocated chromosome was placed on the top of the respective spike. The green signals on chromosomes represent the *Th. elongatum* chromatin and the red signals indicate the centromere. The chromosome name was placed on the right of the chromosome. The diseased spikes were photographed at 21 d after inoculation with *Fusarium graminearum*.

**Fig. S3.**
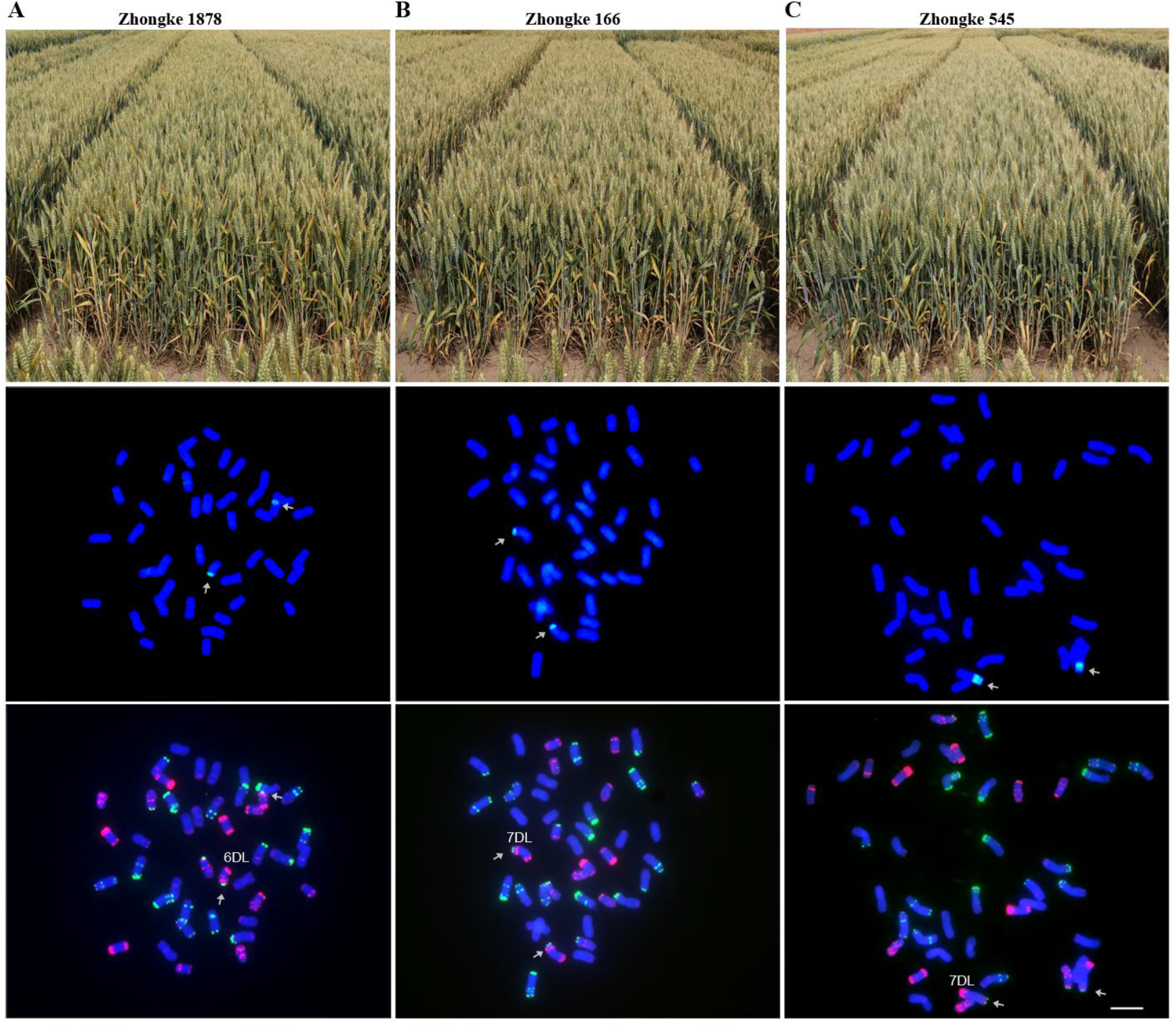
Field plant photograph on yield trial plot and cytological analysis of three translocation lines. Upper panel: Field plant photograph of three translocation lines. Middle panel: FISH analysis on three translocation lines by using 7EL-1 as green probe. The green signal represent *Th. elongatum* chromatin. The arrow indicated the translocated chromosome. Lower panel: FISH analysis on three translocation lines by using pAsⅠ (red) and pSc119.2 (green) as probe. The arrow indicated the translocated chromosome. (A) Field plant photograph on yield trial plot and cytological analysis of Zhongke 1878. (B) Field plant photograph on yield trial plot and cytological analysis of Zhongke 166. (C) Field plant photograph on yield trial plot and cytological analysis of Zhongke 545.

**Fig. S4.**
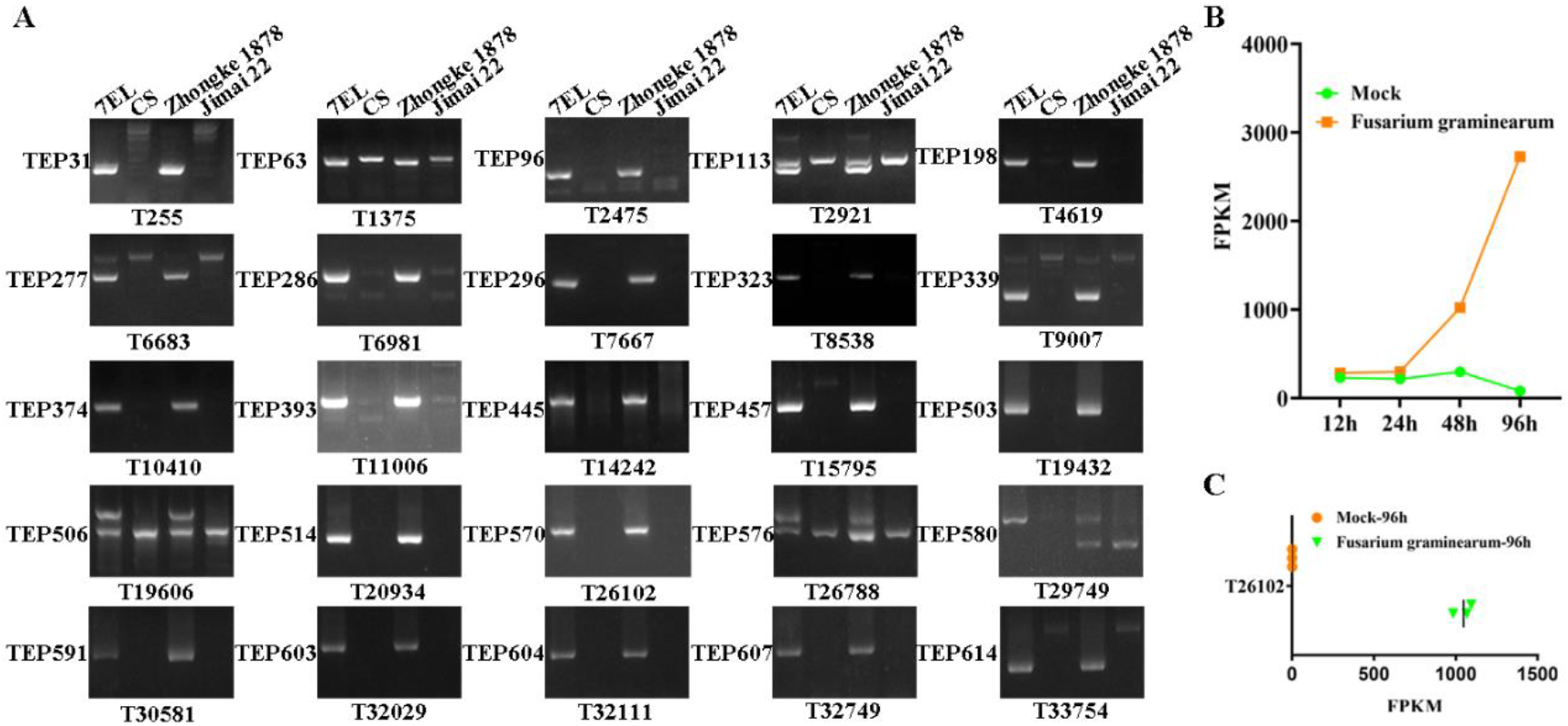
Specific transcripts for alien chromosome and transcriptome analysis on T26102. (A) Twenty-five specific alien transcripts identified by PCR. PCRs were performed by using the genomic DNA of the wheat-*Th. elongatum* addition line 7EL, Chinese Spring (CS), Zhongke 1878 and Jimai 22. The marker name was placed at the left of the electrophoretogram and the transcripts name placed at the bottom of the electrophoretogram. (B) Expression pattern of T26102 in Zhongke 1878 between Mock and *Fusarium graminearum* (Fg) treatment. The Mung bean soup without *Fusarium graminearum* was used as Mock treatment. (C) Expression analysis of T26102 in the addition line 7EL at 96h after Mock and Fg treatment.

**Fig. S5.**
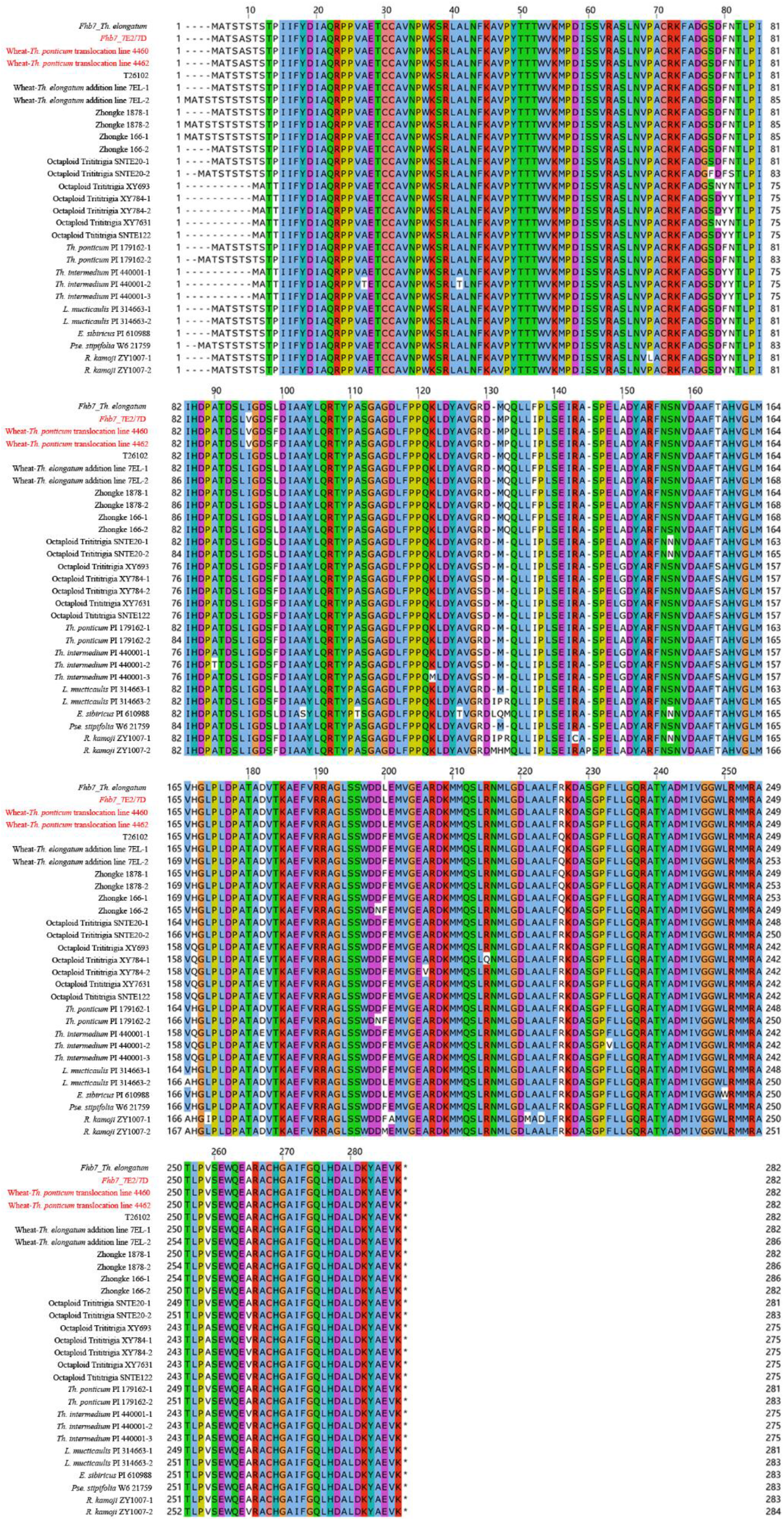
Protein sequence alignments of the *Fhb7* homologs in Triticeae. The sequences used for alignment were derived from five genera in Triticeae, including *Thinopyrum*, *Elymus*, *Leymus*, *Pseudoroegneria* and *Roegneria*. Fhb7_*Th. elongatum*: the GST-encoding Fhb7 candidate cloned from diploid *Th. elongatum*. Fhb7_7E2/7D: the GST-encoding Fhb7 candidate cloned from *Th. ponticum* 7E2 chromosome from the 7E2/7D substitution lines on Chinese Spring background. The protein sequences of T26102 homologs from wheat-*Th. ponticum* translocation lines 4460 and 4462 were identical to Fhb7 in the 7E2/7D substitution line.

**Fig. S6.**
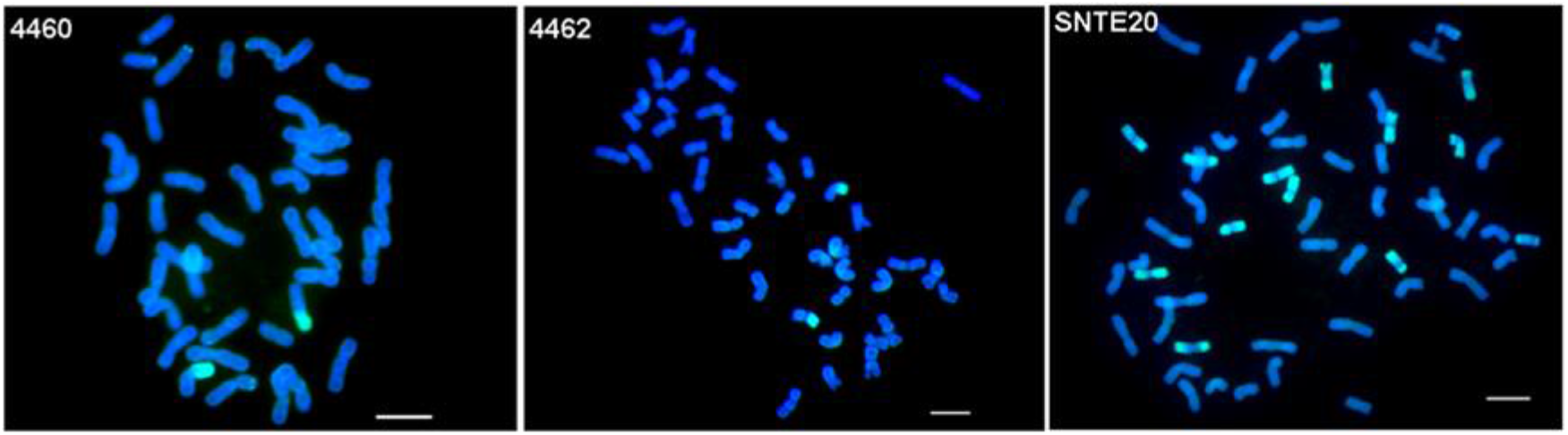
Cytological analysis of wheat-*Thinopyrum* derivate carrying *Fhb7* homolog. 4460 and 4462, wheat-*Th. ponticum* translocation lines; SNTE20, octaploid partial amphiploids developed from *Th. ponticum*.

**Fig. S7.**
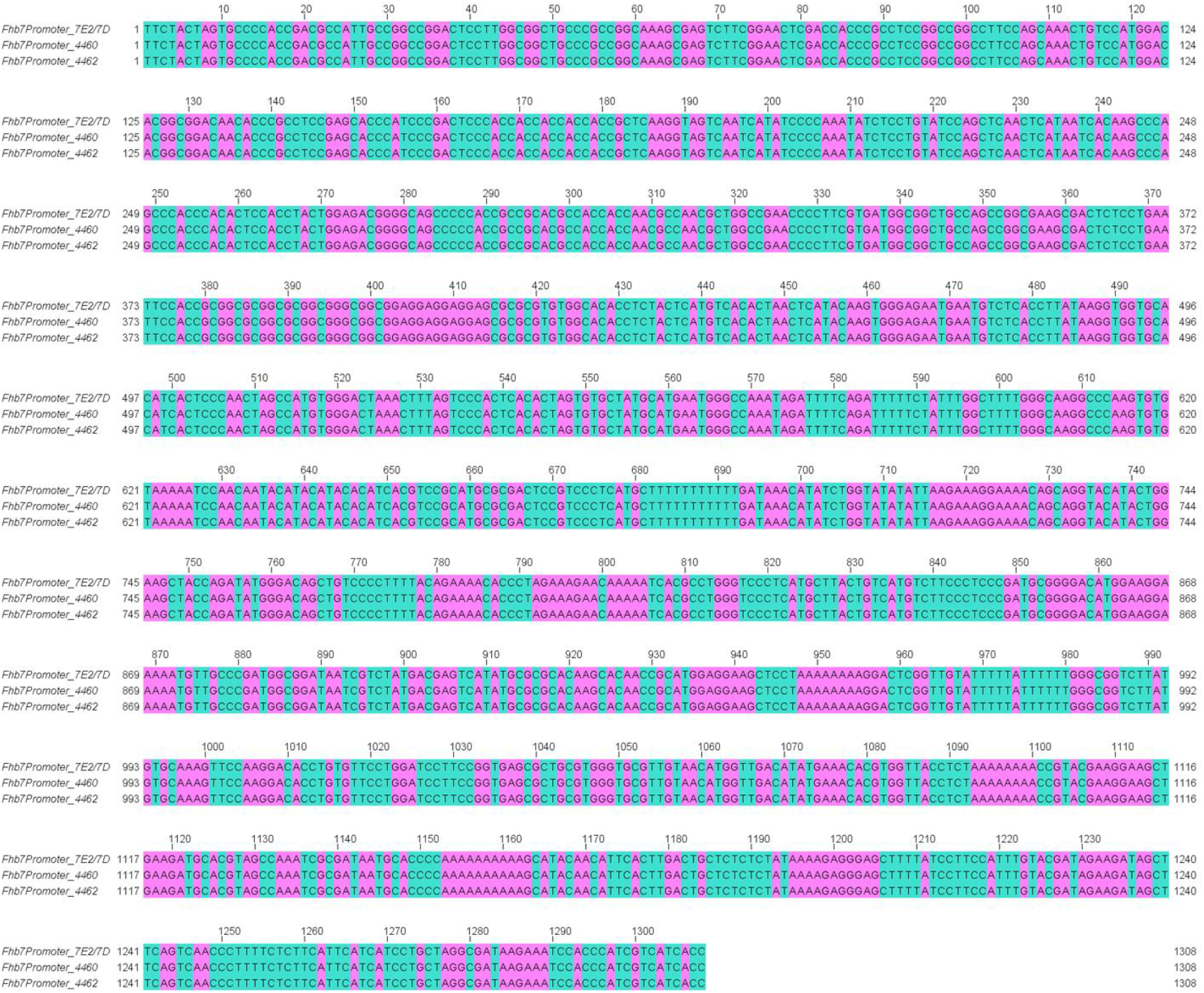
Promoter sequence alignment of the *Fhb7* homologs. *Fhb7*Promoter_7E2: the promoter sequence of the GST-encoding *Fhb7* candidate was from the *Th. ponticum* chromosome 7E2 of the 7E2/7D substitution line on Chinese Spring background.

**Fig. S8.**
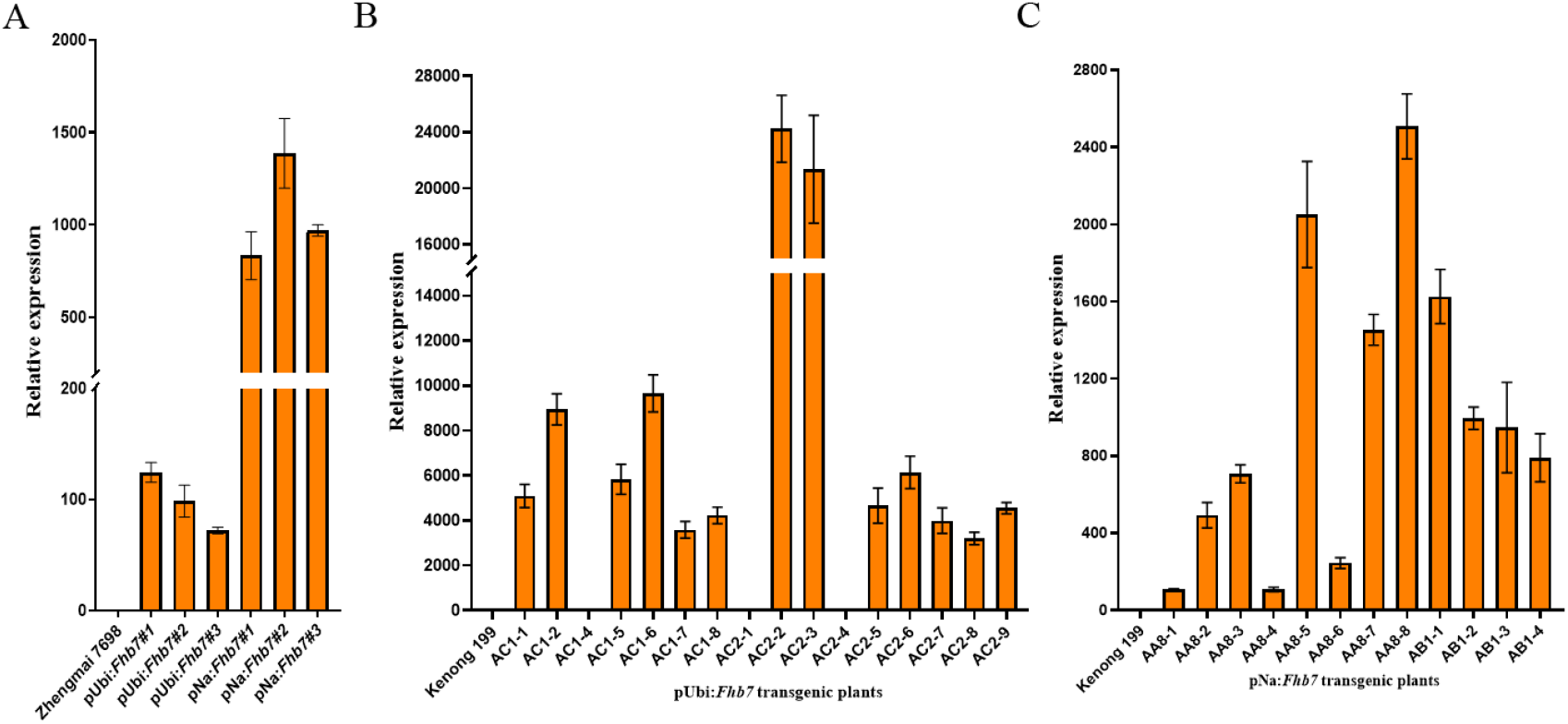
Analysis of the *Fhb7* homolog expression in transgenic wheat plants. (A) Analysis of T26102 expression in transgenic wheat plants on the background 19AS161, Jimai 22 and Zhongmai 175. (B) Analysis of the *Fhb7* expression in transgenic wheat plants on the background Zhengmai 7698. pUbi:*Fhb7* indicated that *Fhb7* was driven by the *ubiquitin* promoter. pNa:*Fhb7* indicated that *Fhb7* was driven by the native promoter. (C) Analysis of the *Fhb7* expression driven by the *ubiquitin* promoter in T0 transgenic wheat plants on the background Kenong 199. AC1and AC2 represented two transgenic lines. (D) Analysis of the *Fhb7* expression driven by the native promoter in T1 transgenic wheat plants on the background Kenong 199. AA8 and AB1 represented two transgenic lines.

**Supplementary Data 1** The sequences of twenty-five specific transcripts in line Zhongke 1878.

**Supplementary Data 2** Distribution of the *Fhb7* homologs in Triticeae.

## Notes

### Competing Interest Statement

The authors have declared no competing interest.

